# Computational design of pH-sensitive binders

**DOI:** 10.1101/2025.09.29.678932

**Authors:** Green Ahn, Brian Coventry, Ella Haefner, Shayan Sadre, Jenny Hu, Mimosa Van, Buwei Huang, Isaac Sappington, Adam J. Broerman, Mauriz A. Lichtenstein, Matthias Glögl, Inna Goreshnik, Dionne Vafeados, David Baker

**Affiliations:** Institute for Protein Design, University of Washington, Seattle, WA, USA; Department of Biochemistry, University of Washington, Seattle, WA, USA; Howard Hughes Medical Institute, Seattle, WA, USA; Molecular and Cellular Biology Program, University of Washington, Seattle, WA, USA; Department of Molecular and Cell Biology, University of California at Berkeley, Berkeley, CA 94720, USA; Department of Biochemistry and Molecular Biology, Hamilton College, Clinton, NY, USA; Xaira Therapeutics, South San Francisco, CA, USA; Graduate Program in Biological Physics, Structure and Design, University of Washington, Seattle, WA, USA; Department of Chemical Engineering, University of Washington, Seattle, WA, USA; Institute for Chemistry and Biochemistry, Freie Universität Berlin, Berlin, Germany

**Author notes:** Correspondence to (D.B.) and (G.A.). These authors contributed equally and are listed in alphabetical order (Green Ahn, Brian Coventry, Ella Haefner, Shayan Sadre).

## Abstract

pH gradients are central to physiology, from vesicle acidification to the acidic tumor microenvironment. While therapeutics have been developed to exploit these pH changes to modulate activity across different physiological environments, current approaches for generating pH-dependent binders, such as combinatorial histidine scanning and display-based selections, are largely empirical and often labor-intensive. Here we describe two complementary principles and associated computational methods for designing pH-dependent binders: (i) introducing histidine residues adjacent to positively charged residues at binder-target interfaces to induce electrostatic repulsion and weaken binding at low pH, and (ii) introducing buried histidine-containing charged hydrogen-bonding networks in the binder core such that the protein is destabilized under acidic conditions. Using these methods, we designed binders that dissociate at acidic pH against ephrin type-A receptor 2, tumor necrosis factor receptor 2, interleukin-6, proprotein convertase subtilisin/kexin type 9, and the interleukin-2 mimic Neo2. Fusions of the designs to pH-independent binders of lysosomal trafficking receptors function as catalytic degraders, inducing target degradation at substoichiometric levels. Our methods should be broadly useful for designing pH-sensitive protein therapeutics.

## Main Text

Dynamic pH gradients are fundamental to eukaryotic cell biology, driving intracellular trafficking, lysosomal activity, and tumor progression. Within cells, compartments such as endosomes and lysosomes are acidified to regulate internalization, degradation, and enzymatic activity^1,2^. In the extracellular milieu, pathological settings such as the tumor microenvironment (TME) are characterized by a lowered pH^3,4^. In nature, pH-sensitivity is often driven by specific hydrogen-bond networks or electrostatic interactions. Interaction between IgG and neonatal Fc receptor (FcRn) is strengthened at lower pH due to the formation of a salt bridge between protonated histidine and glutamate^5^. In Chikungunya virus glycoproteins, hydrogen bonding is weakened at lower pH as protonated histidine can no longer accept a hydrogen bond from serine^6^. Electrostatic repulsion between histidine and charged residues such as histidines, lysines, and arginines weakens binding in several influenza A hemagglutinin complexes^7,8^ (**Fig. S1**). Many biologics and drug modalities have exploited pH changes for maximal efficacy. For example, antibody dissociation from antigen in endosomes paired with recycling via FcRn enables multi-turnover capture of antigens, resulting in longer lasting efficacy and lower dose of treatment^9,10^. Catalytic extracellular degraders, such as catalytic lysosome targeting chimeras (cataLYTACs)^11^, CYpHERs^12^, or sweepers^13^, depend on pH-dependent dissociation of the targets to achieve catalytic degradation of proteins. Moreover, conditionally activated therapeutics, such as antibodies and checkpoint inhibitors, have been developed to switch “on” in the acidic TME, thereby enhancing therapeutic windows^14–19^ (**Fig. 1A**). Despite the numerous applications of pH dependent binders, there are few computational approaches for generating these–most work has utilized random histidine mutagenesis combined with display-based selection to generate pH-dependent binders^11,20,21^, but this process is often laborious and lacks mechanistic clarity for design principles (**Fig. 1B**).

**Figure 1.**
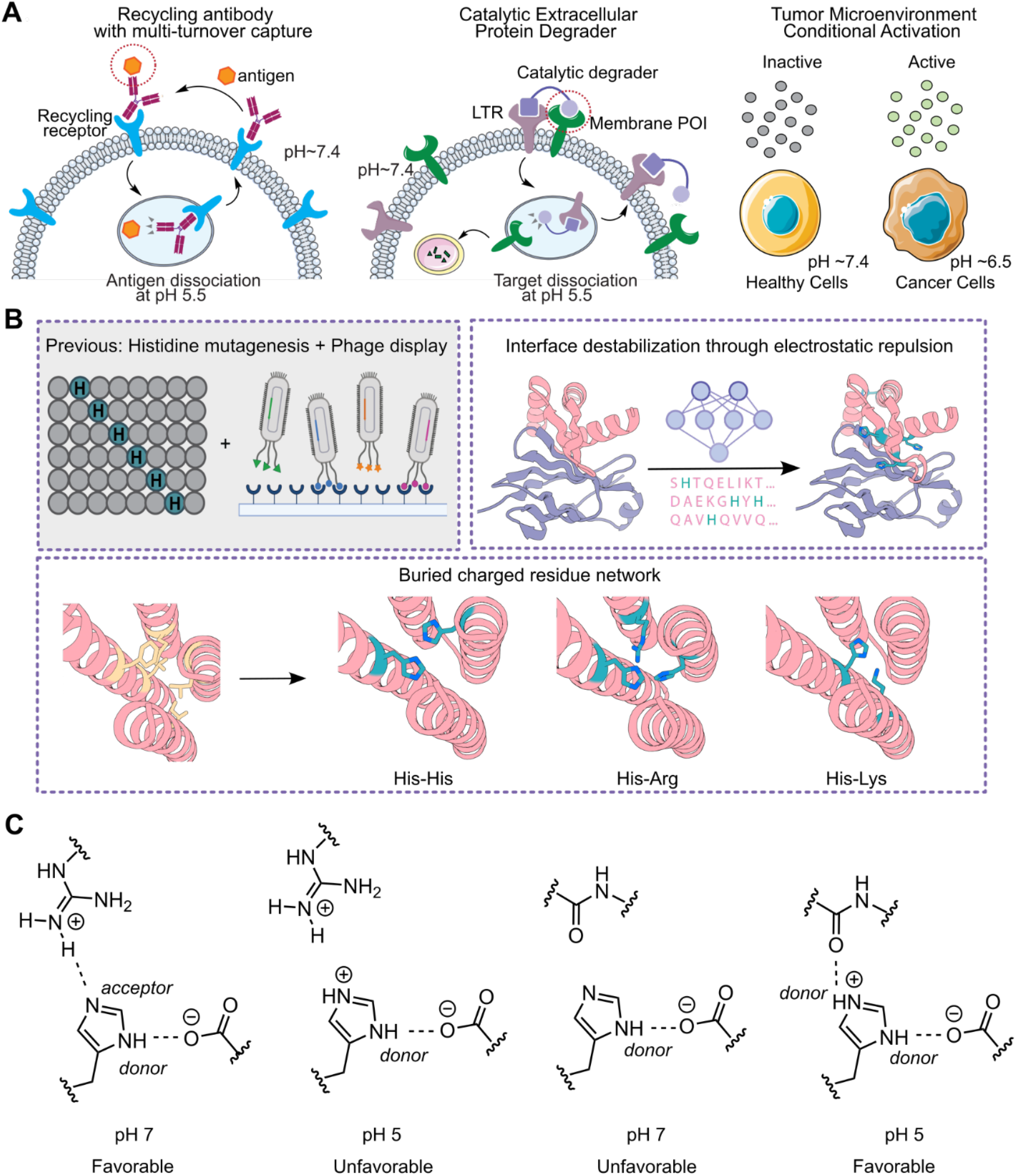
Overview of pH-sensitive binders. **A**. Applications of pH-sensitive binders for recycling antibody with multi-turnover capture, catalytic extracellular protein degrader, and conditional activation in the tumor microenvironment. pH-sensitive interactions are highlighted with red circles. LTR=lysosome trafficking receptor. **B**. Strategies to generate pH-dependent binders. Previously, histidine mutagenesis and phage display have been commonly used. In this manuscript, we developed two directed computational design methods: interface destabilization, incorporating electrostatic repulsion at the interface, and monomer destabilization through buried charged residue networks in the binder core. **C**. Histidine acts as a hydrogen bond acceptor and donor at neutral pH; the acceptor bond is disrupted at pH 5, weakening the interaction (left two panels). Histidine acts as a single hydrogen bond donor at neutral pH and double hydrogen bond donor at acidic pH, strengthening the interaction (right two panels).

We reasoned that recent advances in de novo protein design could be extended to enable the computational design of pH-dependent binders. Deep-learning-based protein design methods such as RFdiffusion^22^ and ProteinMPNN^23^ have enabled the generation of stable binders with high affinity, specificity, and tunable properties. We set out to explore combining physics-based and deep-learning-based computational methods to encode pH-sensitivity into designed binders. We focused on two overall strategies: first, making the protein-protein interface between the designed binder and target pH-sensitive, and second, making the folding of the monomeric binder pH-dependent (**Fig. 1b**).

## Results

### Defining molecular features for pH-sensitivity

Histidine is the only residue with a sidechain pKa near neutral pH, making it particularly sensitive to biological pH gradients. The pH in circulation is ∼7.4, in the endosome, 5.0-6.0, and in the TME, 6.0-6.5^1^. Histidine can both accept and donate hydrogen bonds with charged residues depending on its protonation state^24^. When protonated at pH below the pKa (of ∼6.5), histidine can act as a donor for two hydrogen bonds but not as an acceptor^25,26^, whereas when unprotonated at pH above the pKa, it can act as a donor in one hydrogen bond and an acceptor in a second hydrogen bond; this second hydrogen bond is disrupted upon transfer to acidic pH 6 (**Fig. 1C**). This property has been exploited to design pH-dependent protein homo-oligomers containing nine hydrogen bonds in which histidine acts as a donor and an acceptor–while stable at pH 7, when the pH is reduced to 5, there is a highly cooperative unfolding transition driven by the unfavorable interaction between the newly protonated histidine and the hydrogen bond donating sidechain^27,28^.

Designing “one sided” pH-dependent binders to targets of interest is a harder challenge than “two sided” design of pH-dependent oligomers as only one side of the interface can be changed. To investigate how changes in histidine donor or acceptor state impact the pH dependence of binding affinity in the “one sided” case, we first took an unbiased approach by increasing the number of histidine residues on designed binders and evaluating their pH-sensitivity. We used RFdiffusion for backbone generation and deep learning-based sequence design, ProteinMPNN, with global histidine bias at varying levels to generate designed binders against the oncogenic receptor ephrin type-A receptor 2 (EphA2) and proprotein convertase subtilisin/kexin type 9 (PCSK9). This initial approach yielded binders with increased number of histidine residues, but little pH-sensitivity (**Fig. 2A**). We concluded that specifying the hydrogen bond donor or acceptor status of histidine residues in designs, rather than simply increasing the overall number of histidines, would be necessary to program pH-dependent changes in affinity.

**Figure 2:**
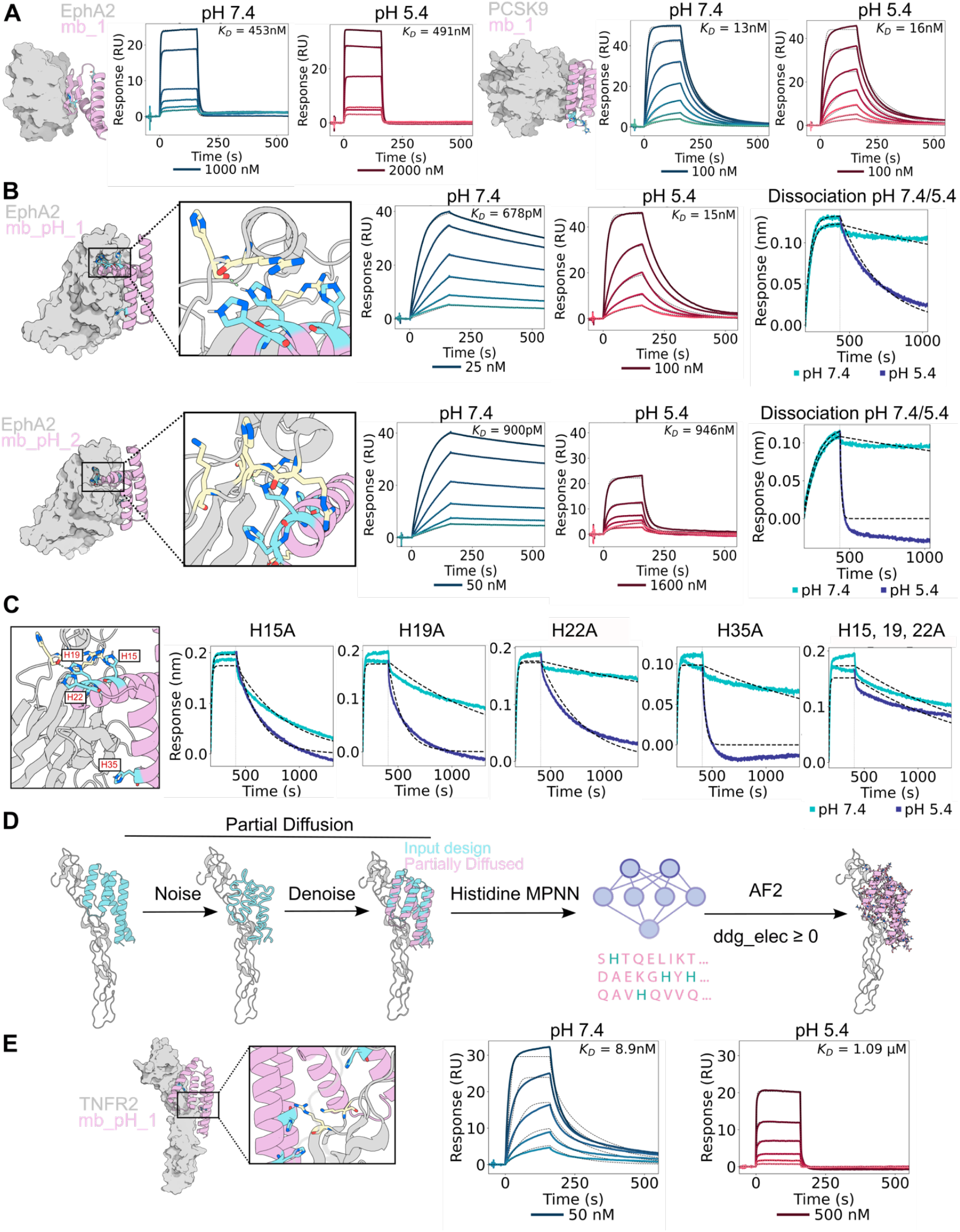
Design of pH-sensitive binder-target interface. **A**. Surface Plasmon Resonance (SPR) curves of EphA2 and PCSK9 binders with histidine-bias ProteinMPNN at pH 7.4 and 5.5. **B**. Design models of pH-sensitive EphA2 binders following yeast display screening. Binding affinities were determined by SPR at pH 7.4 and 5.4 (first two plots). Association at pH 7.4 and dissociation at 5.5 showed faster dissociation at pH 5.5 (third plot). **C**. Evaluation of histidine mutations from EphA2_pH_1 binder. Dissociation rates were determined through BLI at pH 7.4 and 5.4. **D**. Design pipeline for pH-sensitive binders incorporating electrostatic interaction considerations. **E**. Design model and SPR curves of pH-sensitive TNFR2 binder at pH 7.4 and 5.4.

### Designing pH-sensitivity through incorporation of histidine-positive charge interactions at interfaces

We reasoned that specifically focusing on designing hydrogen bond networks across the binder-target interface in which histidines act as acceptors would result in faster dissociation at lower pH. Protonation of the imidazole side chain in acidic pH would disrupt its ability to accept hydrogen bonds, leading to destabilization of the interaction with the donating sidechain. We used partial diffusion^29^ of two previously designed binders against EphA2 to generate a large library of related backbones, and histidine-bias ProteinMPNN to add histidines at the binder-target interface. In a first attempt to quantify the effects of the change in hydrogen bonding patterns at low and high pH, we used Rosetta to sample histidines in neutral and protonated forms, and determined the number of interface hydrogen bonds in which histidine acts as an acceptor, the number in which it acts as a donor, and the degree of burial of each hydrogen bond. As buried hydrogen bonds are expected to have a greater effect on stability and binding affinity, we formed a “pH score” by weighting the difference in the number of acceptor and donor hydrogen bonds by the solvent accessibility (see Methods for details). Using the pH score as a filter, we selected designs containing buried histidines acting as hydrogen bond acceptors and screened 12,000 designs on yeast at pH 5.4 and pH 7.4 to generate a large dataset to identify potential parameters that correlated with decreased affinity at low pH (**Fig. S2**).

We identified four pH-sensitive designs **(Fig. 2B and Fig. S3**) which displayed up to 1000-fold weaker binding at pH 5.4 compared to 7.4 (**Fig. 2B**). To simulate the effect of the drop in pH in the endosome following binding in the extracellular milieu, we carried out bilayer interferometry (BLI) experiments with association at pH 7.4 and dissociation at pH 5.4, and found that the four designs dissociated much faster at pH 5.4 (**Fig. 2B**, right panel). All four pH-sensitive designs showed histidines on the binder interface in close proximity to positively charged arginines on the target, in some cases without direct hydrogen bonding interactions (**Fig. 2B**) Mutational analyses of the histidines in design EphA2_pH_1 revealed that the pH-sensitivity is most conferred by residues His15 and His19 (**Fig. 2C**). Inspection of the design models showed that His15, His19, and His22 were placed near arginine residues on the target protein creating electrostatic repulsion (3.7Å, 7.6Å, and 4.9Å respectively between imidazole nitrogens at the ε2 position of histidines and the terminal guanidium nitrogen of arginines) (**Fig. 2C**). We investigated the factors that differentiated the pH-dependent binders from the non-pH dependent ones more broadly over the 12,000 designs in the yeast display library, and found that the designs with weaker affinity at lower pH had an increase in (unfavorable) electrostatic binding energy (ddG_elec; computed using Rosetta using a distance dependent dielectric) at pH 5 versus pH 7 **(Fig. S4**). Thus, electrostatic repulsion should be considered in designing pH-sensitivity in addition to direct hydrogen bonding.

We next explored whether we could generate pH-sensitive binders with a higher success rate by incorporating the above insights. After incorporating pH-sensitive histidines into existing binders at the binder-target interface by using partial diffusion to generate closely related backbone structures and ProteinMPNN with histidine bias to place histidines on the binder, we selected for designs with a predicted increase in electrostatic repulsion at low pH (ddg elec ≥ 0) and at least one histidine accepting a hydrogen bond from a positively charged residue (**Fig. 2D**). We applied this method to a designed TNFR2 binder^29^, and experimentally characterized 43 designs. 42 of the 43 expressed, and 6 of the binders displayed greater than 2-fold weaker binding at pH 5.4 (**Fig.S5**). The most pH-sensitive binder displayed 122-fold weaker binding at pH 5.4 (**Fig. 2E**). Analysis of the combined EphA2 and TNFR2 datasets suggested that electrostatic repulsion is the major contributor to pH-sensitivity: all of the pH dependent binders in both design sets contained two or more contacts between protonated histidine residues and surrounding cationic residues, including histidine, lysine, or arginine. pH-sensitive EphA2 binder showed up to 11 histidine-cationic interactions, and pH-sensitive TNFR2 binder had up to 8 of such interactions (**Fig. S6**).

### Designing pH-sensitivity through monomer destabilization

Although our interface histidine design pipeline can generate pH-sensitive binders, it modifies the binder interface and relies on the presence of positively charged residues on the target. For designed binders that have strong binding affinity and cross-species reactivity, backbone remodeling or complete sequence redesign of the interface may be undesirable. To preserve these properties, we developed a complementary strategy to introduce a buried network of histidine with positively charged residues into the binder core. Upon protonation of histidines, electrostatic repulsion between the now positively charged histidine and the second cationic residue would destabilize the binder and induce faster dissociation. We used a PyRosetta script as described in the Methods to install histidines that hydrogen bond to lysine, arginine, or histidine in the core of designed binders, and where possible incorporated an additional hydrogen bonding residues to create three residue networks, focusing on more deeply buried regions (see Methods). ProteinMPNN was used to redesign the region around the mutations, and we selected designs that had AlphaFold2 (AF2)^30^ predicted alignment error (pae) interaction and interface root mean square deviation (RMSD) values similar to or improved over predictions for the original designed binder-target complex, and with the energy of the primary histidine - histidine, arginine, or lysine hydrogen bond less than -0.5 kcal/mol (**Fig. 3A**). Because this approach relies on buried hydrogen bonding networks rather than extensive interface redesign, we reasoned that it could be broadly applicable across diverse targets without requiring large library screening and would not be limited by a requirement for positively charged residues at the target interface.

**Figure 3:**
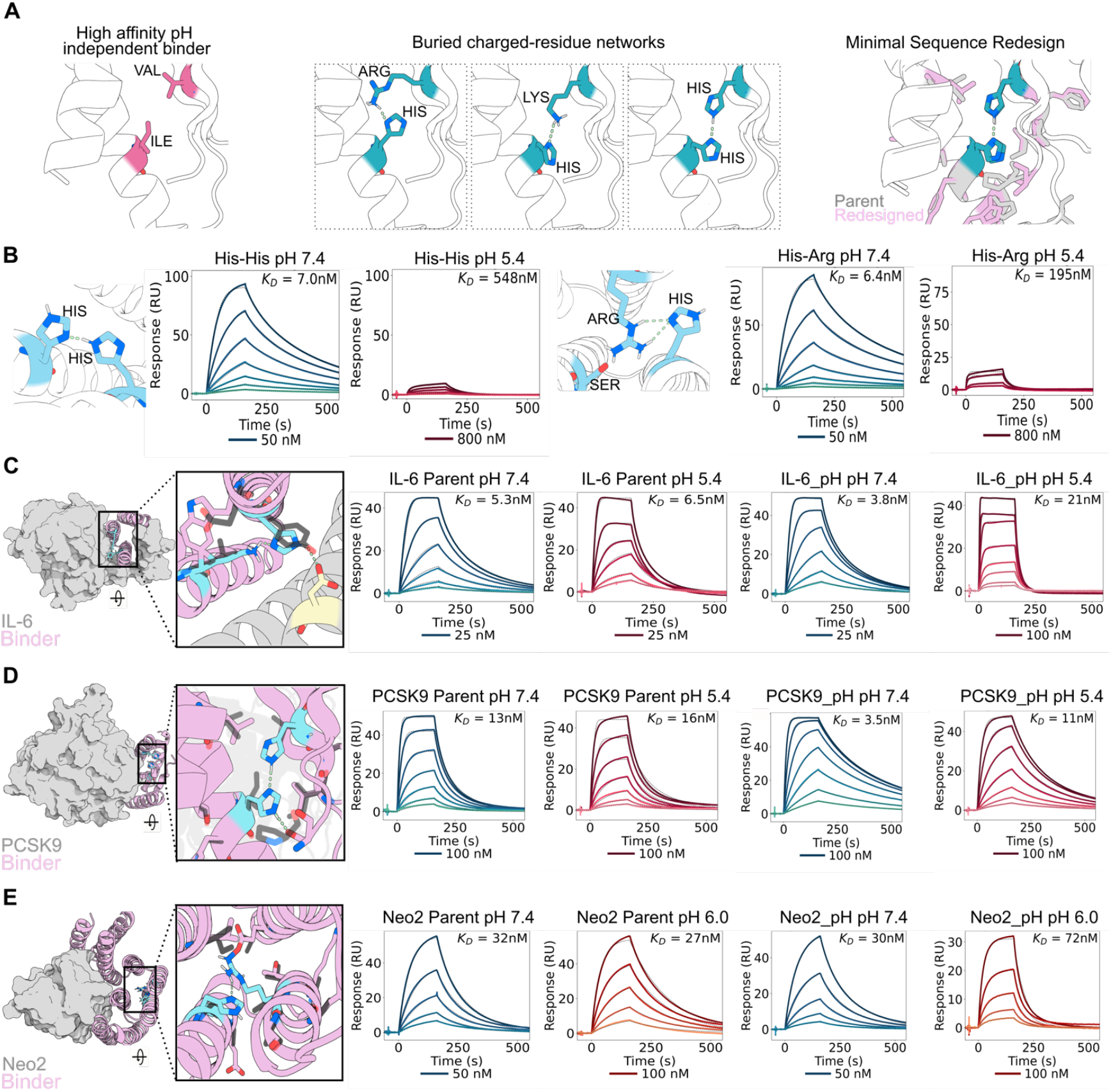
Design of pH-sensitive binders through buried charged residue hydrogen bonding network. **A**. Schematic of the design pipeline. **B**. SPR traces at pH 7.4 and 5.4 for pH-sensitive TNFα binders with His-His or His-Arg network. **C**. IL-6 pH-sensitive binder with hydrogen bond network zoomed in (left-most panel). SPR traces at pH 7.4 and 5.4 for parental binder (IL-6 Parent) and pH-sensitive binder (IL6_pH). **D**. PCSK9 pH-sensitive binder with hydrogen bond network zoomed in (left-most panel). SPR traces at pH 7.4 and 5.4 for parental binder (PCSK9 Parent) and pH-sensitive binder (PCSK9_pH). **E**. Neo2 pH-sensitive binder with hydrogen bond network zoomed in (left-most panel). SPR traces at pH 7.4 and 6.0 for parental binder (Neo2 Parent) and pH-sensitive binder (Neo2_pH). Hydrogen bonds are indicated in blue dashed lines, network residues indicated in cyan, and parental residues indicated in grey.

We sought to apply these design principles to four therapeutically relevant targets for which we had obtained high-affinity de novo binders. We first applied this pipeline to a designed TNFα binder, and found that 48 out of 72 designs retained binding to TNFα, 7 had affinities tighter than 25 nM, and of these, 3 were pH-sensitive. The design models for two of these binders contained buried histidine-histidine and histidine-arginine networks (**Fig 3B**). The top pH-sensitive binder contained a histidine-histidine network and had a 79-fold reduction in binding affinity at pH 5.4 when compared to pH 7.4 (**Fig. 3B**). These designs also showed a faster dissociation rate at pH 5.4 (**Fig.S7A**).

We next generated pH-dependent binders to interleukin-6 (IL-6), PCSK9, and the interleukin-2 mimic Neo2^31^. 36 designs were tested from two different IL-6 binder scaffolds, and 8 designs containing the same network from one backbone were pH-sensitive with the most sensitive having a 6-fold weaker binding at pH 5.4 (**Fig. 3C,. Fig. S8B**). For PCSK9, from 40 binders tested, 32 retained binding affinity less than 25 nM, and of those, 7 were pH-sensitive with the most sensitive having a 3.5-fold weaker binding at pH 5.4 (**Fig. 3D,. Fig.S9A-C**). For Neo2, of 48 binders tested, 33 retained affinity greater than 30 nM. Two of these dissociated faster at pH 6.4 (the approximate pH of the TME) and had greater than 2-fold weaker binding at pH 6.0 (**Fig. 3E,. Fig.S10A**,**B**). Size-exclusion chromatography profiles showed a predominant monomeric peak with minor dimeric species in some instances, consistent with the parent binders for each target (**Fig. S7B, S8C, S9B, S10C**).

Throughout these cases, we identified multiple pH-sensitive variants with networks positioned at different locations within the proteins (**Fig.S9D**). Most networks were buried within the protein core, although in some cases (**Fig.S9C**,**S10B**), networks conferring pH-sensitivity were positioned closer to the binder-target interface. For example, in the IL-6 binder, a histidine-lysine network on the binder lies in close proximity to an asparagine on IL-6 (**Fig. 3C**). Mutational analysis demonstrated that this network was essential for pH-sensitivity, as disrupting the histidine or the lysine abolished pH dependence. Mutation of a proximal tryptophan incorporated during ProteinMPNN redesign reduced binding affinity 6-fold (**Fig. S8A)**. Thus, our method offers advantages over traditional histidine scanning by maintaining high affinity while introducing pH-sensitivity.

### Development of catalytic extracellular degraders

We sought to apply the designed pH-sensitive binders for therapeutic applications. Lysosome targeting chimeras (LYTACs) are heterobifunctional extracellular degraders that engage lysosome trafficking receptor (LTR) and a protein target of interest^32–34^. By recruiting a target protein to an endogenous LTR, LYTACs promote lysosomal degradation of extracellular and membrane proteins. Previously, we developed fully protein-based LYTACs through de novo protein design^35^. Despite the promising degradation results, previous versions of LYTACs dissociated from the LTR in the late endosome, where the pH drops to 5.5, and remained bound to the target protein during lysosomal trafficking. As a result, LYTACs are stoichiometric, where each drug molecule degrades along with the target protein. To ensure recycling of LYTACs with the LTR, the LTR binder must remain bound to the receptor even in the acidic endosome, while the target binder must dissociate from the target (**Fig. 4A**). We designed a pH-insensitive binder (IRigd1) against an LTR, insulin-like growth factor 2 receptor (IGF2R) **(Fig. S11)**. We then fused this binder to either Protein G or pH-sensitive Protein G to assess for catalytic internalization of IgG. We observed that the pH-sensitive Protein G LYTAC showed significantly higher potency and enhanced internalization of IgG even at concentrations below that of the target, indicating substoichiometric degradation (**Fig. 4B**).

**Figure 4:**
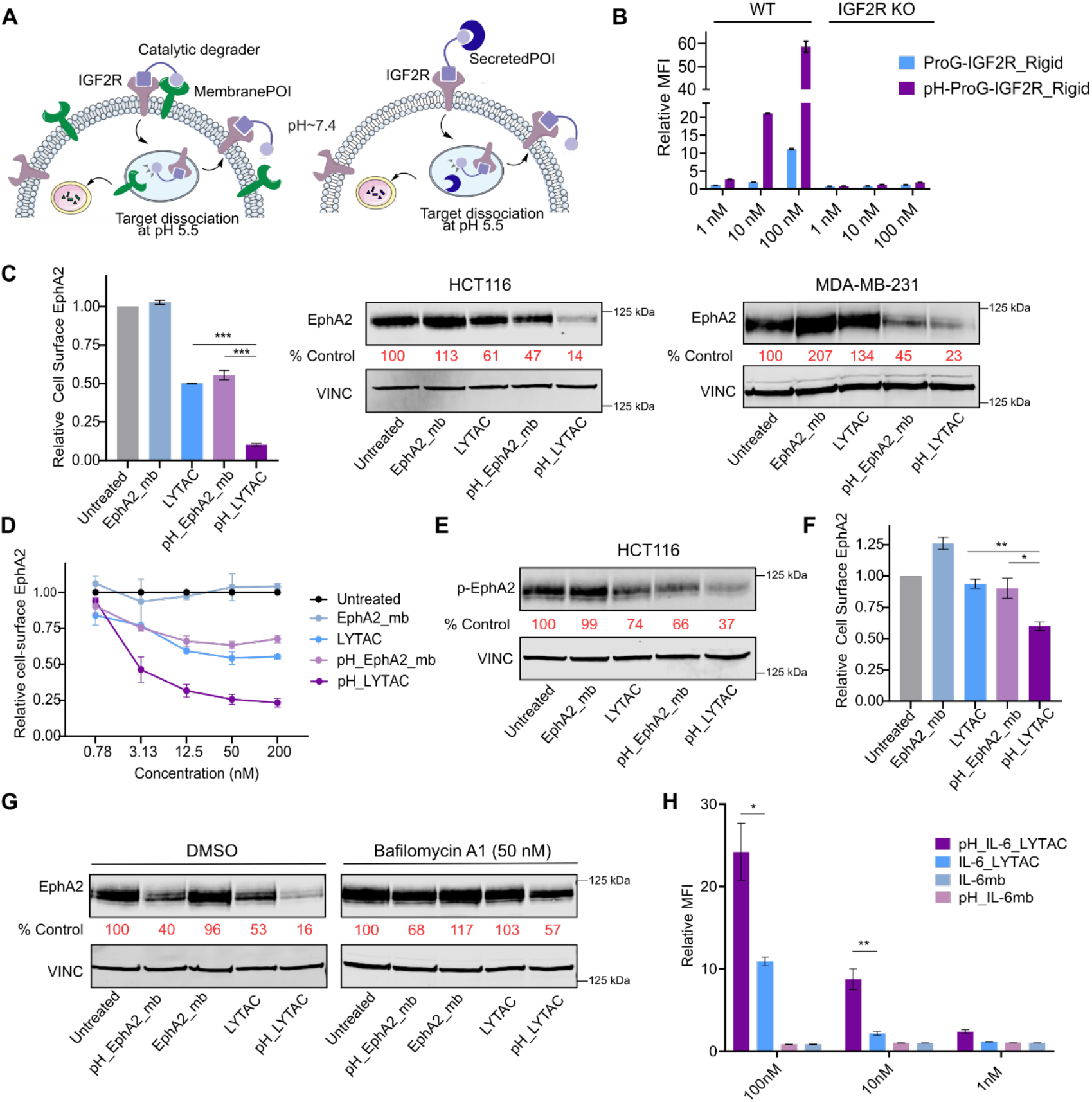
Catalytic degradation of membrane and secreted proteins using designed pH-sensitive binders. **A**. Schematic of catalytic extracellular degradation of membrane protein and secreted protein with designed pH-sensitive binders. **B**. Internalization of 100 nM of AF647-labeled IgG with varying concentrations of Protein G-LYTAC or pH-Protein G-LYTAC. HEK293T cells were treated with LYTACs and IgG for 24 h. Mean fluorescence intensity (MFI) was determined by live cell flow cytometry. **C**. Degradation of cell-surface EphA2 in HCT116 cells as determined by flow cytometry (left). Degradation of total EphA2 in HCT116 (middle) and MDA-MB-231 (right) cells as determined by western blot. Cells were treated with 250 nM designed proteins for 24h. **D**. Dose-response curve for cell-surface EphA2 degradation in HCT116 cells following 24h treatment with designed proteins. **E**. Western blot analysis of phosphorylated EphA2 (p-EphA2, Ser897) following 24h of treatment. **F**. Degradation of cell-surface EphA2 following a washout. HCT116 cells were treated with 250 nM of designed proteins for 24 h, then were washed out of extracellular treatment and incubated with fresh media for 6h. Mean fluorescence intensity (MFI) was determined by live cell flow cytometry. **G**. Western blot degradation of EphA2 in HCT116 cells following co-treatment of DMSO or Bafilomycin A1 (50 nM) with designed proteins. **H**. Internalization of 50 nM biotinylated IL-6 precomplexed with AF647 streptavidin following 24h treatment of LYTACs (10, 1 nM indicate substoichiometric ratios of the degraders) in K562 cells. MFI was determined by live cell flow cytometry. Data in **B, C, D, F, H** are shown as the mean in three independent experiments ± s.e.m. *P* values were determined by unpaired two-tailed *t*-tests. Data in **C, E, G** are representative of three independent replicates.

Catalytic degraders of EphA2 could be useful for cancer therapy, as EphA2 has an oncogenic role in breast cancer, lung cancer, glioblastoma, and melanoma^36^. We fused the pH-sensitive EphA2 minibinder (pH_EphA2_mb) to a pH-insensitive IGF2R minibinder (IRigd1) to test for catalytic degradation. These pH-sensitive LYTACs (pH_LYTACs) induced more potent degradation of EphA2 in two different cancer cell lines, HCT116 and MDA-MB-231, as assessed by cell-surface staining and western blot (**Fig. 4C,. Fig. S12a**), with dose-dependent degradation (**Fig. 4D**). The pH_LYTACs also resulted in the ablation of phosphorylated EphA2 (S897), an oncogenic form of the receptor (**Fig. 4E,. Fig. S11b**). To determine whether the degraders can induce catalytic degradation, we performed a wash-out experiment, removing the LYTAC from solution and assessing whether degradation persisted. We treated cells with LYTACs for 24 hours, then performed stringent washing of the cells and incubated them in fresh media. Although EphA2 levels were partially recovered, pH_LYTACs maintained significant degradation of EphA2, whereas the non-pH-sensitive LYTACs showed complete reversal of degradation (**Fig. 4F,. Fig. 11c**). These results demonstrate that the LYTACs derived from pH-sensitive binders are able to achieve multiturnover through recycling. Inhibition of lysosomal acidification with bafilomycin A1 blocked degradation (**Fig. 4G)**, confirming that degradation goes through the endolysosomal pathway.

Although secreted proteins such as cytokines are common therapeutic targets, their high levels in circulation requires high doses of conventional inhibitors and antibodies. Catalytic extracellular degradation enables degradation of substoichiometric levels of target, potentially increasing the therapeutic index. Because IL-6 is a cytokine that drives chronic inflammation and cytokine storm^37,38^, a catalytic degrader offers a powerful strategy to deplete IL-6 in circulation. To investigate catalytic degradation of IL-6, we employed the pH-sensitive IL-6 binder developed from the buried charged-residue hydrogen-bond network pipeline. Compared to parental IL-6_LYTACs, the pH-IL-6_LYTACs showed higher potency and substoichiometric degradation (**Fig. 4H**).

## Discussion

This study establishes generalizable approaches for designing pH-sensitive binders. We observed that random histidine incorporation rarely weakens binding at low pH, highlighting the need for histidine incorporation guided by considerations of hydrogen bonding and electrostatic repulsion. We developed two complementary design methods that encode pH responsiveness through distinct mechanisms: first, through direct interface destabilization by inducing electrostatic repulsion upon histidine protonation, and second, through destabilization of the monomer structure through incorporation of histidine-containing buried charged networks. The two design strategies are best suited to different use cases. Interface histidine design can be applied to either existing binders or de novo scaffolds and is most effective when the target surface presents exposed cationic residues, while incorporation of buried charged residue networks requires scaffolds with sufficiently large cores. We show that control over pH-sensitivity enables the design of catalytic degraders: fusions of a pH-independent binder of a recycling receptor to a pH-dependent binder of a target of interest mediate the binding of target at the cell surface at neutral pH, the release of the target in the late endosome at low pH, and the recycling of the target-free fusion to the cell surface for additional rounds of degradation.

Our design strategies can be applied not only to de novo designed proteins, but also to natural proteins and antibodies, allowing engineering next-generation therapeutics with tunable pharmacokinetic profiles and tumor selectivity. The mechanistic principles from this study can be applied to generate proteins that bind more tightly at low pH through design of histidines that are hydrogen bond donors instead of acceptors for applications such as facilitating recycling of therapeutics through receptors beyond FcRn and IGF2R, as well as selective binding in the TME. Finer control over pH thresholds - particularly optimizing for better responsiveness around pH 6.5 by shifting the pKa of histidine through tuning of the surrounding residues - will be important for improving the therapeutic window for TME activation. More generally, programmable pH-sensitivity enables building conditional control into protein-based therapeutics, diagnostics, and synthetic biology tools.

## Methods

### Binder design through RF Diffusion and ProteinMPNN

To design de novo binders to EphA2, IL-6, PCSK9, and Neo2, we initially generated 5000-10,000 backbones using RFdiffusion^22^. We targeted binders using “hotspot” residues to a specific site on the target protein. We used AF2 with an initial guess^39^ and target templating to filter designs. This configuration of AF2 runs without a multiple sequence alignment and keeps the target fixed while predicting the de novo binder. Design models that were predicted from AF2 with pae_interaction <20 were subjected to partial diffusion^29^ for backbone optimization, followed by another round of ProteinMPNN and AF2 filtering. After two rounds of partial diffusion and sequence optimization, designs were selected by filtering on pae_interaction <8 and pLDDT >88, and were subjected to FastRelax to obtain Rosetta metrics^40^. The resulting binders were further filtered on ddG and SAP score^41,42^.

### Binder Screening

All screening except designs tested on yeast surface display were screened by SPR (method details described below). Linear gene fragments encoding binder design sequences were cloned into LM627(addgene #191551) using Golden Gate assembly. Golden Gate subcloning reactions of binders were constructed in 96-well PCR plates in 4 µl volume. One µl reaction mixtures were then transformed into 6µl chemically competent expression strain (BL21 (DE3)). After 1-h recovery in 100 µl SOC medium, the transformed cells were distributed equally into 4 96-deep-well plates containing 1mL of auto-induction media (autoclaved TBII media supplemented with kanamycin, 2 mM MgSO4, 1 × 5052). Cells were grown at 37 °C for 24 hours and harvested by centrifugation (15 min at 4,000*g*). Bacteria were lysed for 15 min at 37 °C in 400 μl lysis buffer (1× B-PER (78243, Thermo Fisher Scientific), 25U/mL Benzonase, 0.1mg/mL Lysozyme, and one tablet of Pierce protease inhibitor tablet per 50 ml culture). Lysates were clarified by centrifugation at 4,000*g* for 10 min, before purification on Ni-NTA Agarose (30250, Qiagen; wash buffer (20 mM Tris pH 8.0, 300 mM NaCl, and 25 mM imidazole) and elution buffer (20 mM Tris pH 8.0, 300 mM NaCl, and 400 mM imidazole)). Elutions were filtered through a 0.2 μm filter plate and subsequently injected in AKTA Pure SEC Mini using HBSEP+ running buffer. Fractions were normalized to 4 μM, and SPR was performed using Single cycle kinetics with six four-step dilutions. Designs with affinities lower than 200nM were scaled up for medium scale production and further characterized (described below).

### Protein production and purification

Minibinders were expressed in E.Coli BL21. Briefly, the gene fragments encoding the design sequences were assembled into LM627 vectors via Golden Gate assembly, and transformed into BL21 strain. Cells were grown in 50 ml culture and protein expression was induced by the autoinduction media. After 20h of expression at 37°C, cells were harvested by centrifugation and lysed with 15 ml of lysis buffer (25 mM Tris-HCl, 150 mM NaCl, protease inhibitor tablet (Roche)). Lysed cells were sonicated and subjected to ultracentrifugation at 14,000g for 30 min. Proteins were purified by immobilized metal affinity chromatography (IMAC). All pH-sensitive binders were cleaved by TEV-protease to remove the His-tag following elution. Proteins were further purified by FPLC size exclusion chromatography using Superdex 75 10.300 GL column (GE Healthcare).

### Interface histidine design with electrostatic repulsion filtering

#### ProteinMPNN with histidine bias

EphA2 and TNFR2 binders were partially diffused. We then used ProteinMPNN with histidine bias (0.5, 1.0, 1.5) to generate the sequence with varying numbers of histidine residues. Designs were selected by filtering on pae_interaction <8 and pLDDT >88.

#### Generation of low pH structures

The pH module of Rosetta was used to generate low pH structures. For this study, only doubly-protonated histidines were considered. The histidines were protonated with Rosetta by setting the pH:pH_mode to True, the pH:pH_value to 0 (versus 7), and then repacking all histidines with a scorefunction whose only scoreterm was e_pH set to 100. The result was that all histidines were doubly protonated. These doubly protonated histidines contain a +1 formal charge as well as different hydrogen bonding patterns.

#### Determining the interface histidine hydrogen bonding pattern score (pH score)

In an early attempt to quantify the effects of the change in h-bonding patterns at low pH versus high pH, an ad hoc sum of terms was created.

pH_score = 1.0 * low_pH_surf_D

+ 3.0 * low_pH_surf_DD

+ 2.0 * low_pH_surf_D_bb

+ 6.0 * low_pH_surf_DD_bb

+ 3.0 * low_pH_core_D

+ 9.0 * low_pH_core_DD

+ 6.0 * low_pH_core_D_bb

+ 18.0 * low_pH_core_DD_bb

+ 1.0 * high_pH_surf_A

+ 1.0 * high_pH_surf_D

+ 3.0 * high_pH_surf_AD

+ 2.0 * high_pH_surf_A_bb

+ 2.0 * high_pH_surf_D_bb

+ 6.0 * high_pH_surf_AD_bb

+ 3.0 * high_pH_core_A

+ 3.0 * high_pH_core_D

+ 9.0 * high_pH_core_AD

+ 6.0 * high_pH_core_A_bb

+ 6.0 * high_pH_core_D_bb

+ 18.0 * high_pH_core_AD_bb

Each term represents the number of histidines that fit a specific set of criteria making cross-chain hydrogen bonds. The criteria are in the name of the term where low_pH denotes the pH 5 pose, high_pH denotes the pH 7 pose, surf/core denotes protein surface versus protein core, A/D/AD/DD denote the hydrogen bonding pattern of each histidine with “A” meaning the nitrogen of the HIS sidechain is accepting a H-bond, “D” meaning the hydrogen of the HIS sidechain is donating a hydrogen bond, “AD” denoting HIS that are both accepting and donating, and “DD” denoting a doubly-protonated HIS making two hydrogen bonds. Finally, “bb” indicates that HIS is making an H-bond to the backbone of the other chain.

#### Determining delta electrostatic ddG

The change in the ddG of binding of the binder at the two different pHs can be assessed by calculating the Rosetta ddG of the binder at the two concentrations and taking ddG_at_pH_5 - ddG_at_pH_7. However, we found that a significant portion of this value was noise from Van der Waals forces and not related to the change in pH. To precisely examine the effect of pH, especially in non-hydrogen bonding pairs, we devised the delta electrostatic ddG, or dddG_elec. This value is calculated by repacking the pose with Rosetta at both low and high pH, and then calculating the ddG using only the fa_elec scoreterm. The result is that this value looks at the change in coulomb’s law energy at pH 5 vs pH.

#### Determining the number of histidine-cationic interactions

We used Rosetta to quantify pairwise distances between histidine on the binder and cationic (histidine, lysisine, arginine) residues on the target structure. The script extracts Cα coordinates and constructs all-by-all pairwise distance matrices. Pairs involving histidine residues are filtered based on chain identity and spatial proximity. For each histidine-cationic residue pair, all possible sidechain rotamers were enumerated, and the shortest non-hydrogen distance was recorded.

### Yeast surface display screening

The yeast surface display screening was performed as described^43^. Briefly, genes encoding the designed binder sequences were transformed into EBY-100 yeast strain. The yeast cells were grown in CTUG medium and induced in SGCCA medium. Induced cells were washed with PBSF (PBS + 1% BSA), and incubated with Streptavidin-phycoerythrin (SAPE, Thermo Fisher, 1:100), and anti-Myc fluorescein isothiocyanate (myc-FITC, 1:100) for 30 mins at room temperature. The cells were washed with PBSF and sorted by FACS (Sony SH800) for FITC positive cells (expression sort). Sorted cells were grown, induced, and incubated with 1 μM biotinylated EphA2 (Acros Biosystems), SAPE, and myc-FITC for binding sorting. Cells that were PE and FITC double-positive were sorted. After another round of sorting without avidity (incubation with EphA2 followed by washing and incubation with SAPE), the yeast cells were titrated with biotinylated EphA2 at different concentrations (100 nM, 10 nM, 1 nM) in PBSF buffer with pH 7.4 and 5.5. The cells were subsequently washed and sorted at two different pH’s. Sorted cells from each subpool were lysed and their sequences were determined via next-generation sequencing.

### Dissociation rate measurements through biolayer interferometry

To measure the dissociation, Streptavidin-coated biosensors (Sartorious) were first loaded with biotinylated target proteins at 50 nM concentration, and a response threshold limit was set to 0.6nm. Sensors were washed with pH 7.4 Octet Buffer (McIlvane’s, .05% Tween20, 1% BSA) and pH 5.4 (McIlvane’s, .05% Tween20, 1% BSA) and incubated with a single selected concentration of binders relative to previously determined KD. (IL-6 100nM, TNFα 100nM, EphA2 100 nM, PCSK9 250nM, Neo2 32 nM). To measure the off-rate (*K*off), the biosensors were then dipped back into the Octet buffer. A reference analyte at the respective pH was used to subtract from the baseline, and an interstep correction was performed. The off-rate *K*off was estimated with the Octet Analysis software using a partial local fit.

### Surface Plasmon Resonance

Binding kinetics were analyzed via Surface Plasmon Resonance (SPR) on a Biacore 8K (Cytiva). Binding to various binders was measured by capturing biotinylated targets (Acro Biosystems: PCSK9 #H82E7,IL-6 #H82Q9,TNFα #H82E3, EPHA2 #H82E4, Neo2 (expressed and purified from previous work^31^), Sino Biological: TNFR2 #10417-H27H-B) using Biotin CAPture Kit (Cytiva #28920234) respectively, by injecting a previously determined concentration of target at a flow rate of 10 µL/min in either pH 5.4 SPR Buffer (McIlvane’s, .05% Tween20) or pH 7.4 (McIlvane’s, .05% Tween20) aiming for a capture level between 250-400 response units. Analytes were diluted in SPR Buffer at the respective pH and injected at a flow rate of 30 µL/min to monitor association. SPR Buffer at the respective pH was used as a running buffer during dissociation at a flow of 30 µL/min. Parallel kinetics was performed using different concentrations in the eight channels. Association/dissociation times and concentration ranges were varied to suit the respective analytes. Binding kinetics were determined by global fitting of curves to kon and koff, assuming a 1:1 Langmuir interaction, using the Cytiva evaluation software.

### Buried charged residue network

#### Pyrosetta Script to Install Histidines

A python script using PyRosetta was created in order to search through designed binders to find pairs of positions to install HIS-HIS H-bonds (later expanded to HIS-ARG/LYS/HIS). The script is loosely based on the tools that underlie MonteCarlo HBNet^44^. RotamerSets for HIS (and potentially ARG/LYS/HIS) are generated for all positions in the binder. These are passed to core.pack.hbonds.hbond_graph_from_partial_rotsets() which generates a map of all h-bonds between all positions. For each pair of positions that can accept a HIS-HIS hydrogen bond (H-bond), the rotamer pair with the best H-bond energy (according to Rosetta) is kept (for each of ARG/LYS/HIS). All pairs of H-bonding rotamers are output as separate PDBs with metrics describing the quality of the H-bond, the depth of the atoms, and the local position of the residues (e.g. core/boundary/surface/interface). In order to quantify the burial of H-bonds, the depth of the H-bond below the molecular surface was calculated. This was performed by using the AtomicDepth method in Rosetta to examine the depth below the molecular surface of both of the atoms participating in the H-bond. The H-bond_depth is then the average depth below the molecular surface of these two atoms.

#### Installing backup histidines

With a script nearly identical to that used to install the initial histidines, a second script was created to produce networks of 3 HIS. This second script used the same functionality to search for all H-bonding rotamers to the first 2 HIS installed and output any rotamer that made a H-bond to them as well as the Rosetta energy of that H-bond.

#### Designing sequence around the histidines

In order to ensure that the proteins still folded and that the HIS pairs actually made their desired H-bond, Rosetta FastDesign biased by ProteinMPNN was used to design the local area around the HIS. The MPNN weights were used to remove amino acids from consideration by Rosetta by calculating the log conditional probabilities and (--conditional_probs_only) and disallowing anything with a log-probability worse than -2.5. From here, two strategies were employed. In the first strategy, all amino acids within 5, 5.5, or 6Å (3 separate trajectories) of the HIS-pair (atom-atom distance) were allowed to redesign. The original amino acids were given energetic boosts of -1, -2, or -3 kcal/mol (3 separate trajectories of each of the 3 distance cutoffs) in order to keep the sequence closer to the starting point. To enable tighter packing, a second strategy was also employed where for each position with an atom-atom distance to the HIS-pair of less than 6Å, all MPNN-allowed amino acids were forced onto that sequence position and the remaining 6Å window redesigned.

#### Evaluation following structural prediction

We used Rosetta scoring to ensure that designed histidine networks remained geometrically consistent and energetically favorable following AF2 structure prediction. For each structure, we computed per-pair energies relative to glycine-substituted baseline and used Rosetta to calculate the hydrogen-bond energy of the histidine network.

### EphA2 degradation via flow cytometry

HCT116 or MDA-MB-231 cells were plated 1 day before the treatment. Cells were incubated with indicated concentration of binders or LYTACs for indicated times. Cells were then washed 3 times with DPBS and trypsinized for <5 minutes and transferred to a 96-well U-bottom plate. Cells were washed twice with FACS buffer (DPBS + 0.2% bovine serum albumin), stained on ice with AF647-anti-EphA2 antibody for 30 min, and washed 3 times with FACS buffer. After the final wash, cells were incubated with Sytox Green and analyzed on the Attune Cytometer (Thermo Fischer).

### Degradation via western blot

HCT116 or MDA-MB-231 cells were plated 1 day before the treatment. Cells were incubated with indicated amount of binders or LYTACs for 24 h. Following the treatment, cells were washed 3 times with DPBS and lysed with RIPA Lysis Buffer supplemented with protease inhibitor, phosphatase inhibitor, and benzonase on ice for 30 min. Cells were scraped and spun down at 21,000g for 15 minutes at 4°C. The supernatant was collected and the concentrations were normalized through a BCA assay. 45 μg of lysates were loaded onto 4-12% Bis-Tris Gel and separated by SDS-PAGE. The gel was then transferred to a nitrocellulose membrane, and blocked with Intercept Blocking Buffer (LICOR) for 1h at room temperature. The blots were stained with primary antibody overnight at 4°C, washed 3 times with TBS-T, and stained with secondary antibody for 1 h at room temperature. LICOR Odyssey CLx Imager was used to image and quantify the blot.

### Internalization of IL-6

K562 cells were plated at 50,000 cells per well of a 96-well plate in 100 μL final volume. LYTAC molecules at 300 nM were precomplexed with 150 nM biotinylated IL-6 and 150 nM Alexa Fluor™ 647 Streptavidin for 1 hour at room temperature. 50 μL of the precomplexed reagents were added to the cells, resulting in a final concentration of 100 nM LYTAC, 50 nM IL-6, and 50 nM AF647 Streptavidin. Serial dilutions were performed for 10 nM and 1 nM concentration mixtures prior to addition. Following 24-hour treatment, cells were washed 3 times with FACS Buffer, stained with Sytox Green, and analyzed on the Attune Cytometer.

## Acknowledgements

We thank K. VanWormer and L. Goldschmidt for technical support, S. R. Gerben for protein production support, and X. Li for LC-MS support. We thank M. Kennedy for manuscript editing. This work was supported by the Audacious Project at the Institute for Protein Design (G.A., I.S., A.J.B.), Jane Coffin Childs Memorial Fund for Medical Research (G.A.), a gift from Microsoft (A.J.B.), Nordstrom Barrier Institute for Protein Design Directors Fund (B.H.), Howard Hughes Medical Institute (D.B., G.A., B.C., I.G., D.V.), Open Philanthropy Project Improving Protein Design Fund, Department of the Defense Defense Threat Reduction Agency grant HDTRA1-21-1-0007 (M.G.), National Institutes of Health’s National Institute on Aging grant R01AG063845 (B.H., I.G., D.B.), Gates Foundation grant INV-043758 (G.A., E.H., S.S., I.S., D.B.), and National Science Foundation 2244288 (M.V.). Figures 1 and 4 were modified from Servier Medical Art, licensed under a Creative Commons Attribution 3.0 Generic License https://smart.servier.com and figure 1 was modified from BioRender.

## Conflict of interest

The authors declare the following competing interests: G.A., B.C., E.H., S.S., and D.B. are in the process of filing a provisional patent application that incorporates discoveries described in this article.

## Contribution

G.A., B.C., E.H., S.S., and D.B. conceived the project. G.A., E.H., S.S., J.H., M.V., B.H. carried out the experiments and interpreted data. G.A. designed initial EphA2 minibinders. E.H. designed pH-sensitive EphA2 minibinders. S.S. and I.S. designed PCSK9 and IL-6 minibinders. M.G. performed partial diffusion on TNFR2 binder. M.A.L. designed the initial minibinder against Neo2. B.C. and A.J.B. developed the concept of buried histidine network. B.C. developed computational methods for calculating Rosetta metrics and generating buried histidine networks. G.A. and D.B. provided supervision. G.A., E.H., S.S., and D.B. wrote the manuscript with input from all authors.

