## Supplementary Information for "Computational design of pH-sensitive binders"

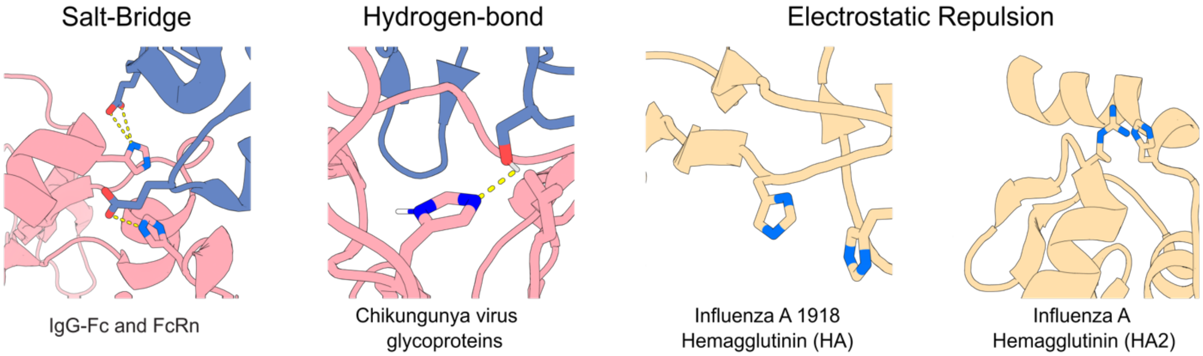


**Fig S1. pH-dependent interactions found in native proteins.** Salt-bridge is indicated in IgG-Fc and FcRn interaction[5] (PDB 1I1A). Hydrogen-bond network is shown in Chikungunya virus glycoproteins[6] (PDB 3N43). Electrostatic repulsion between histidine-histidine or histidine-arginine is shown for influenza A hemagglutinin complexes[7,8] (PDB 4FNK, 1RUY).

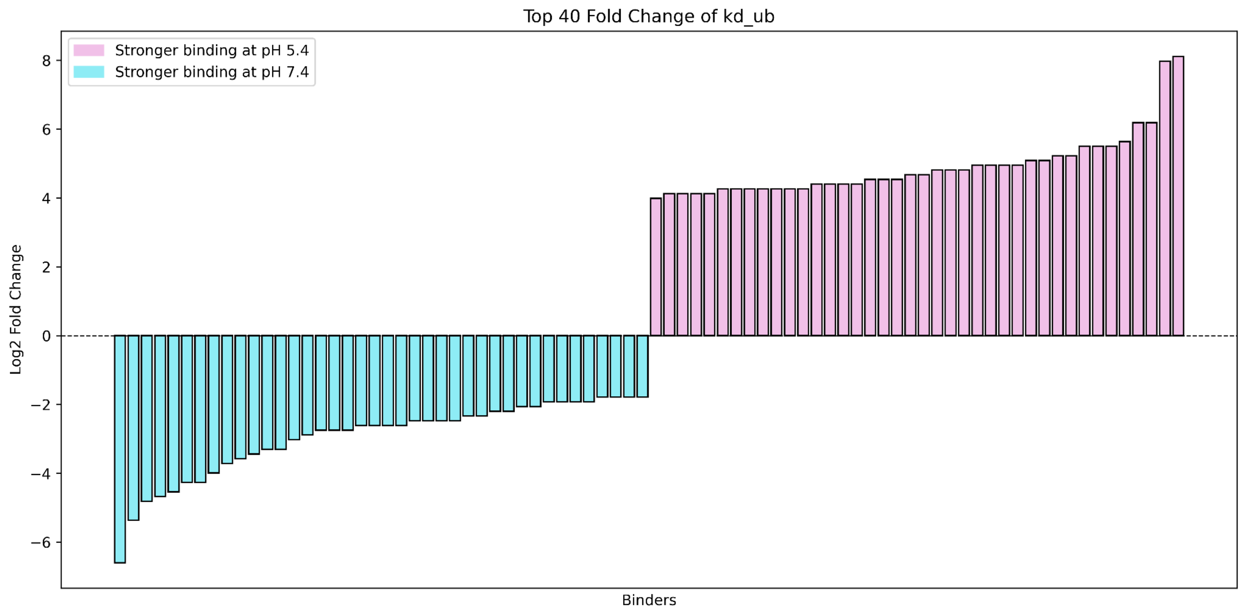


**Fig. S2**. **Yeast log fold change at pH 7.4 vs 5.4 for designed EphA2 binders.**

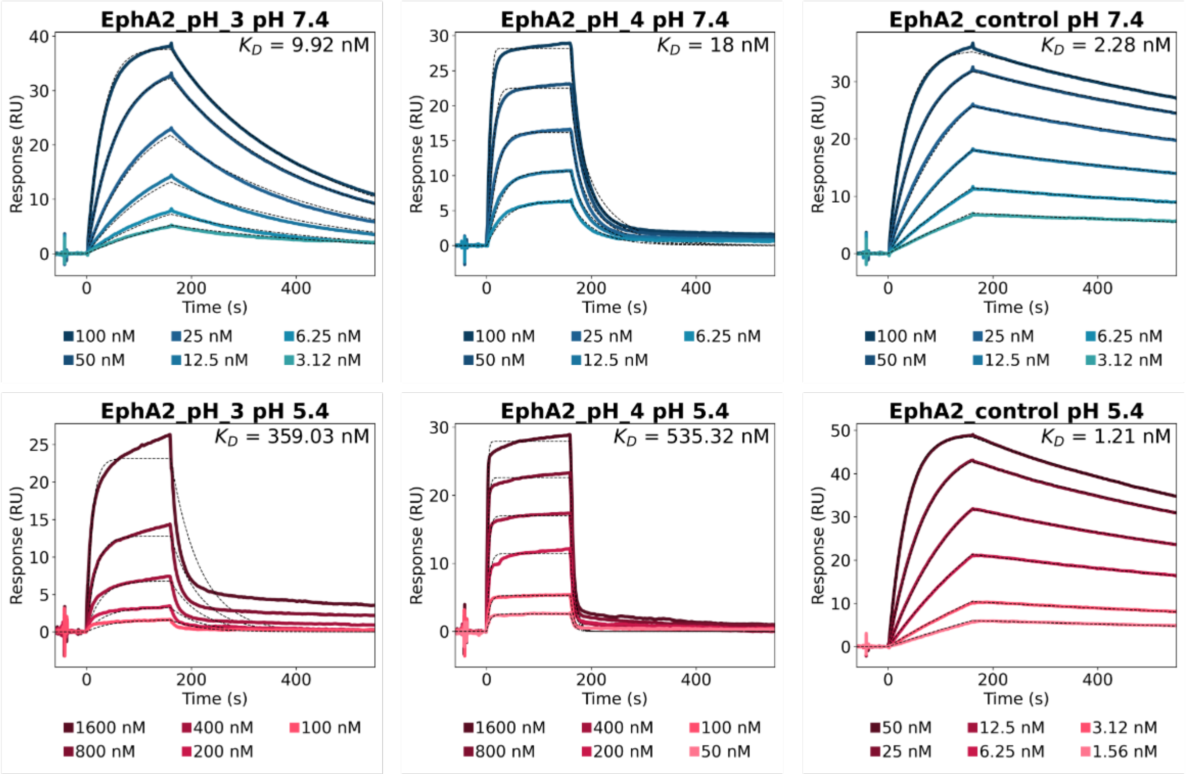


**Fig S3. pH sensitive EphA2 binders.** Binding affinities of designed pH-sensitive EphA2 binders_3, 4, control were determined by SPR at pH 7.4 and 5.4. EphA2_control was a pH insensitive binder utilized in future cell experiments.

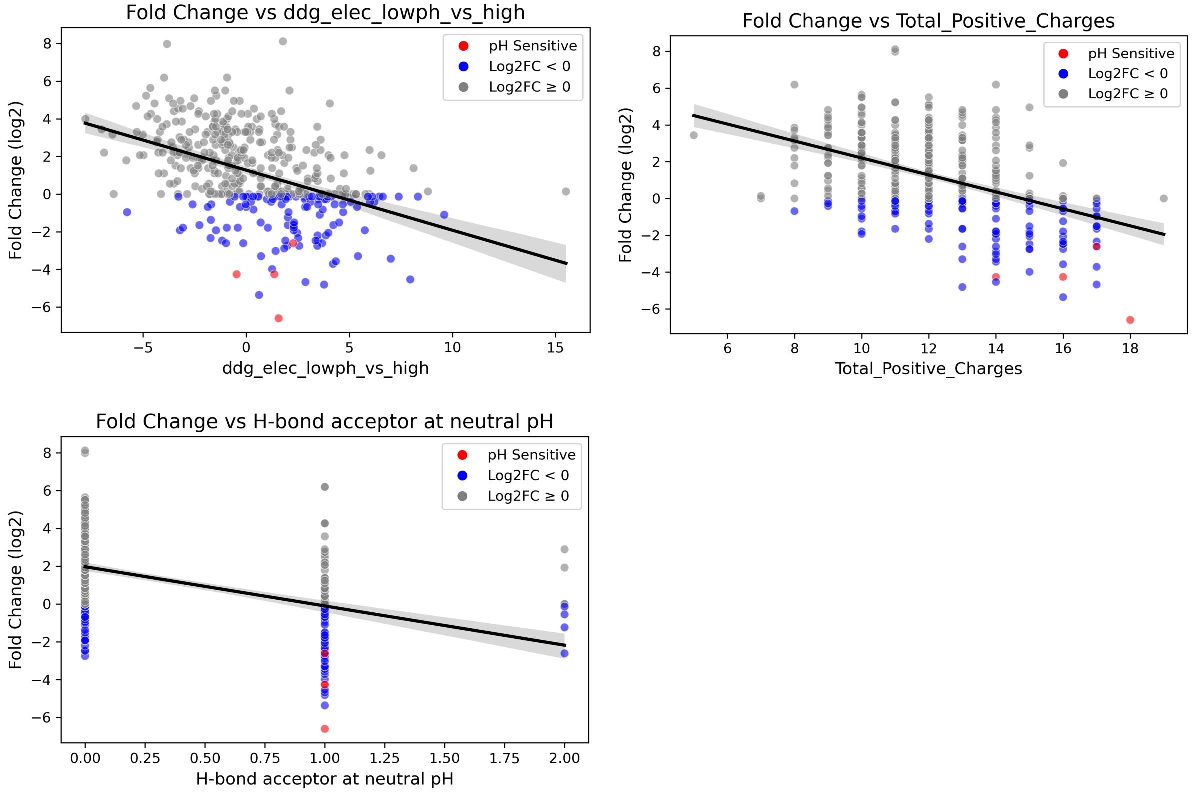


**Fig S4.** Metrics correlating with pH sensitivity. ddg_elec, positive charges, and H-bond acceptor at neutral pH were compared to the log2 fold change of binding from the EphA2 yeast library.

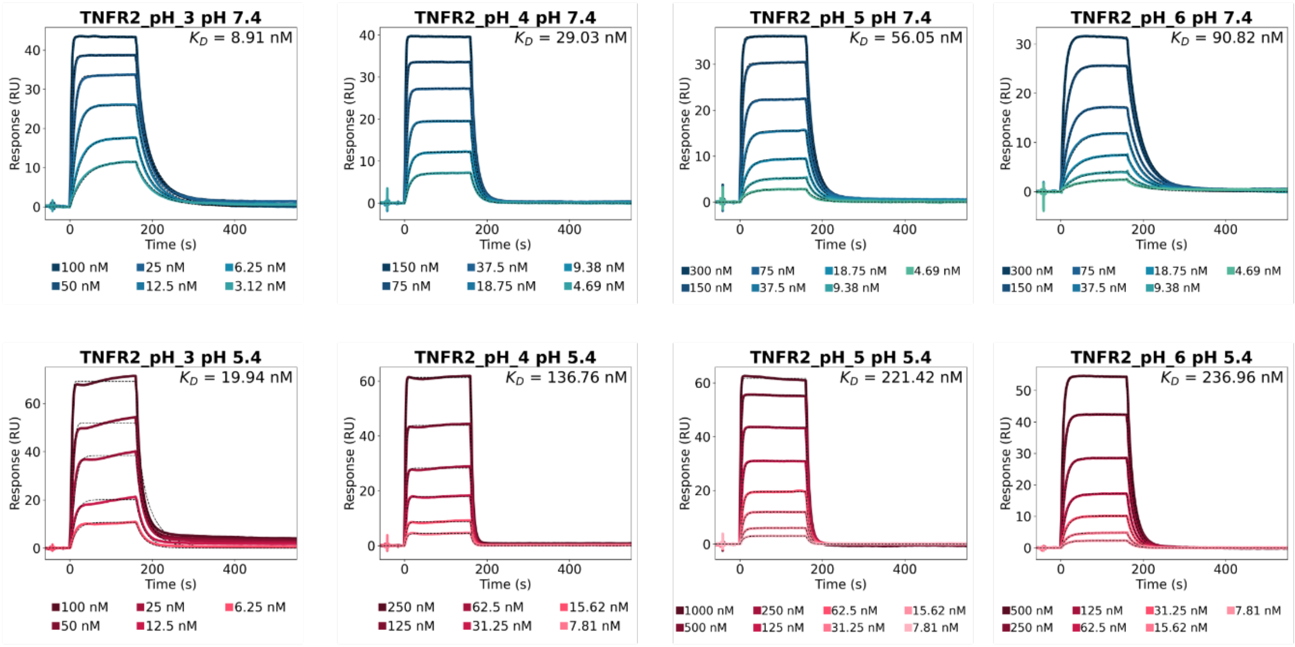


**Fig S5. pH sensitive TNFR2 binders.** Binding affinities of pH-sensitive TNFR2 binders_3, 4, 5, 6 were determined by SPR at pH 7.4 and 5.4.

A

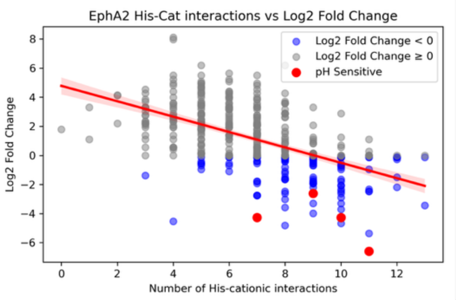


B

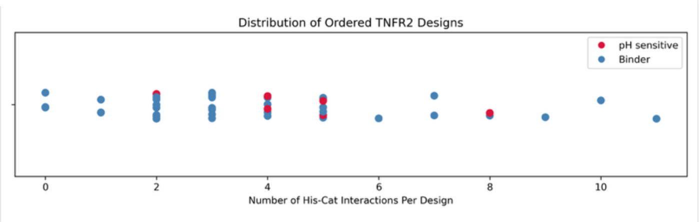


C

**EphA2_pH_1 Binder Interactions**

| binder_residue | distance | target_residue |
|---|---|---|
| 15H | 3.7 | 142R |
| 19H | 7.8 | 199H |
| 19H | 7.6 | 198R |
| 19H | 8.6 | 201K |
| 19H | 8.8 | 142R |
| 22H | 8.7 | 199H |
| 22H | 4.9 | 198R |

**EphA2_pH_2 Binder Interactions**

| binder_residue | distance | target_residue |
|---|---|---|
| 19H | 7.6 | 199H |
| 19H | 5.3 | 198R |
| 19H | 8.2 | 201K |
| 19H | 8.9 | 142R |
| 20H | 8.8 | 198R |
| 20H | 9.8 | 142R |

**EphA2_pH_3 Binder Interactions**

| binder_residue | distance | target_residue |
|---|---|---|
| 19H | 7.6 | 199H |
| 19H | 7.3 | 198R |
| 19H | 8.4 | 201K |
| 19H | 8.9 | 142R |
| 9H | 9.1 | 142R |

**EphA2_pH_4 Binder Interactions**

| binder_residue | distance | target_residue |
|---|---|---|
| 19H | 7.7 | 199H |
| 19H | 7.0 | 198R |
| 19H | 8.3 | 201K |
| 19H | 8.9 | 142R |

**EphA2_control Binder Interactions**

| binder_residue | distance | target_residue |
|---|---|---|
| 25H | 5.7 | 115R |
| 27H | 10.0 | 105K |
| 55H | 5.9 | 116R |

**Fig S6. Calculated electrostatic (histidine-cationic) interactions across designed binders**. **A.** Scatter plot comparing the log2 fold change of binding from the EphA2 yeast library with calculated his-cat interactions. pH-sensitive binders are colored in red. **B.** Distribution of ordered TNFR2 designs and the number of his-cat interactions. The SPR validated pH sensitive binders are colored in red. **C.** Distance in Å between indicated designed binder residue and target residue in 4 different pH-sensitive EphA2 binders and the pH-insensitive control binder. Distances were measured between non-carbon sidechain heavy atoms (HIS – ND1, NE2, ARG – NE, NH1, NH2, LYS – NZ).

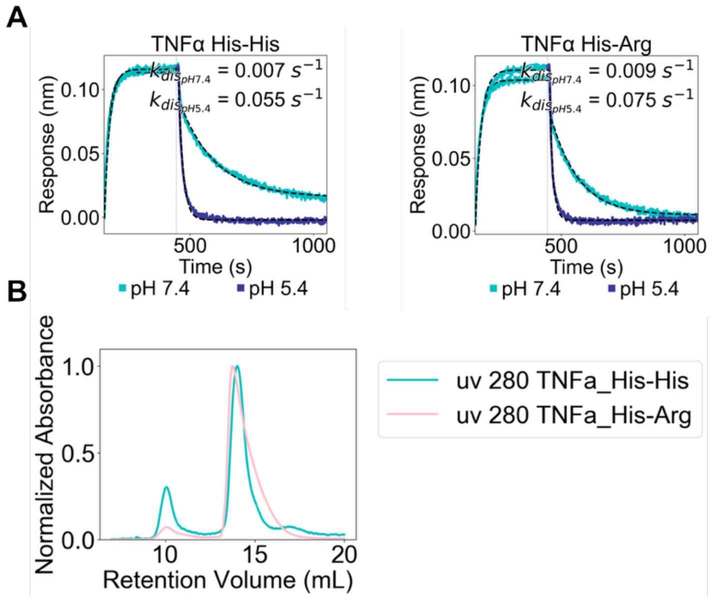


**Fig S7. Characterizations of pH-sensitive TNFα binders. A.** BLI curves of TNFα binders with His-His network or His-Arg network. Association at pH 7.4 and dissociation at 5.4 showed faster dissociation at pH 5.4. **B.** Size exclusion chromatography traces of pH-sensitive TNFα binders.

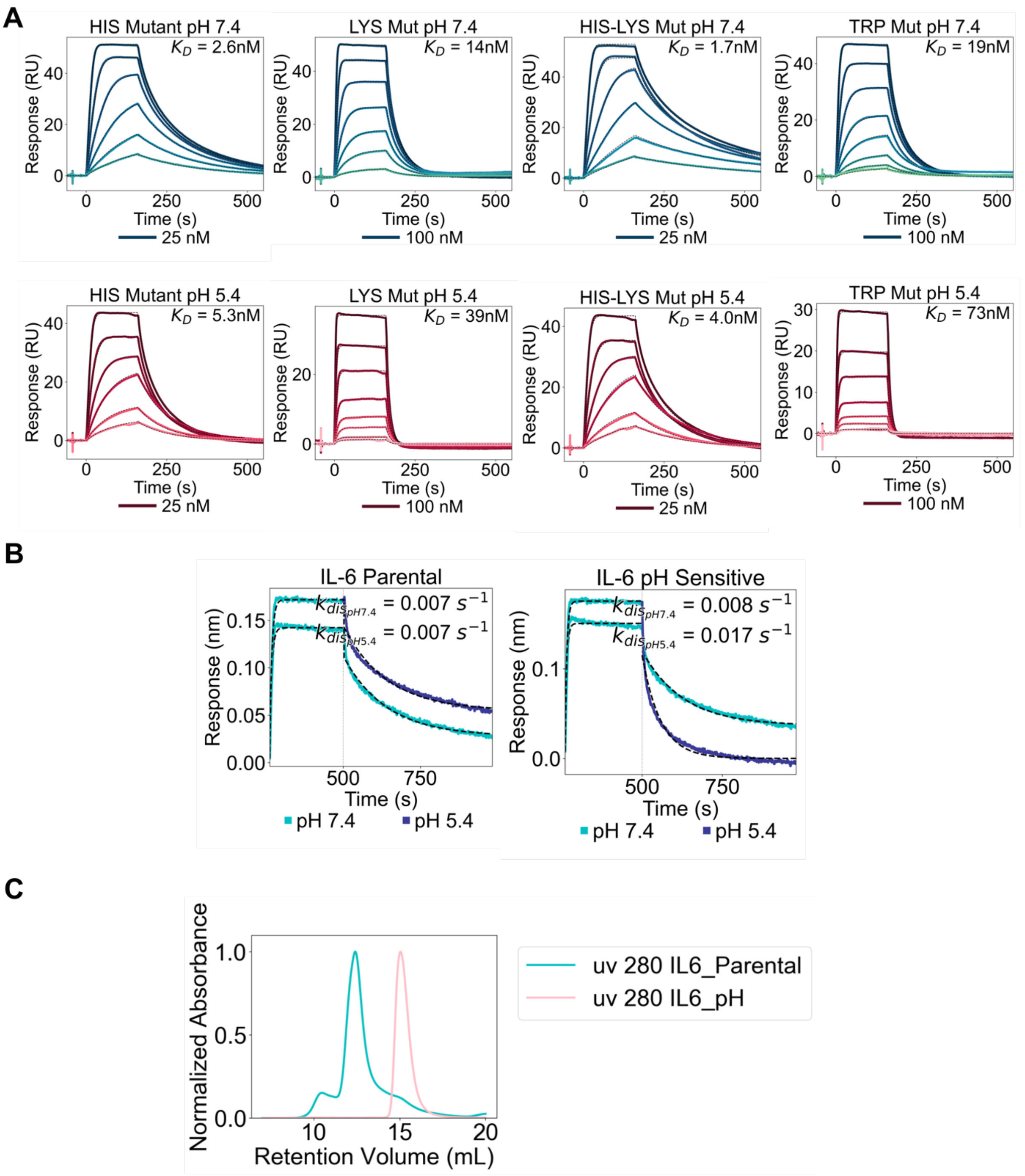


**Fig S8. Characterizations of pH-sensitive IL-6 binders. A.** SPR curves of pH sensitive IL-6 binder variants with the indicated residues restored to the parental sequence at pH 7.4 and 5.4. **B.** BLI curves of parental and pH-sensitive IL-6 binders. Association at pH 7.4 and dissociation at 5.4 showed faster dissociation at pH 5.4 for the pH sensitive binder. **C.** Size exclusion chromatography traces of the parental and pH-sensitive IL-6 binders.

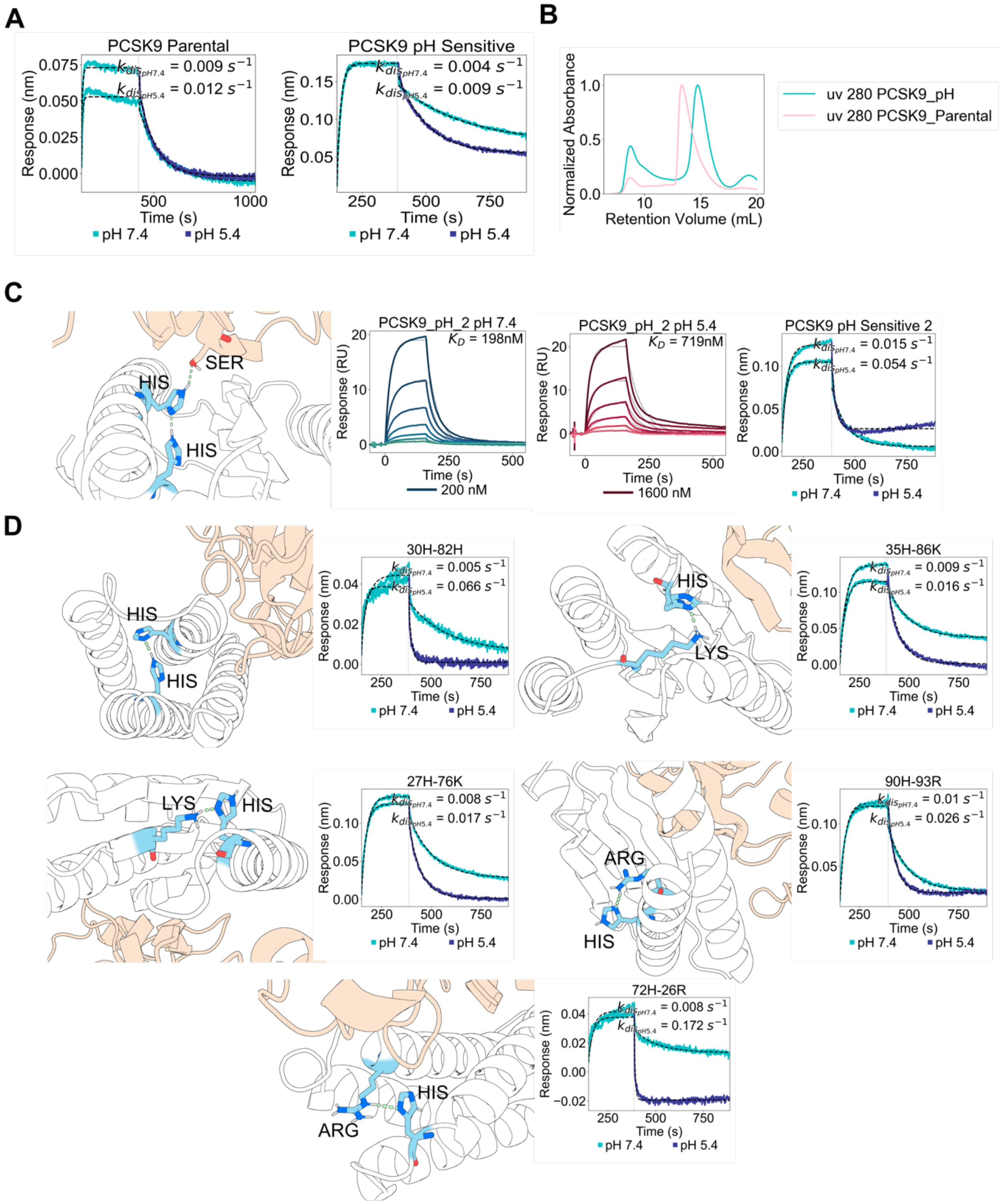


**Fig. S9. Characterizations of pH-sensitive PCSK9 binders. A.** BLI curves of parental and pH-sensitive PCSK9 binders. Association was conducted at pH 7.4 and dissociation at pH 5.4. **B.** Size exclusion chromatography traces of pH-sensitive PCSK9 binders. **C.** Design model and SPR curves of pH sensitive PCSK9 binder_2 at pH 7.4 and 5.4. BLI curves of association at pH 7.4 and dissociation at 5.4 showed faster dissociation at pH 5.4. **D.** Design model and BLI curves of pH-sensitive PCSK9 binders with various charged networks. Association at pH 7.4 and dissociation at 5.4 showed faster dissociation at pH 5.4.

A

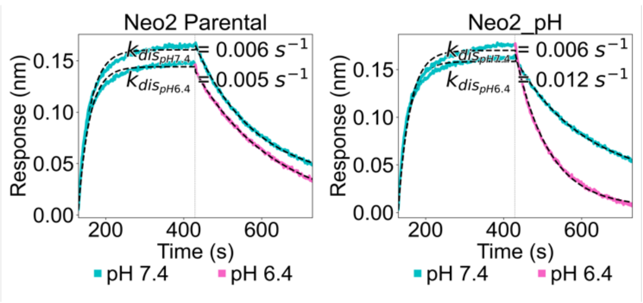


B

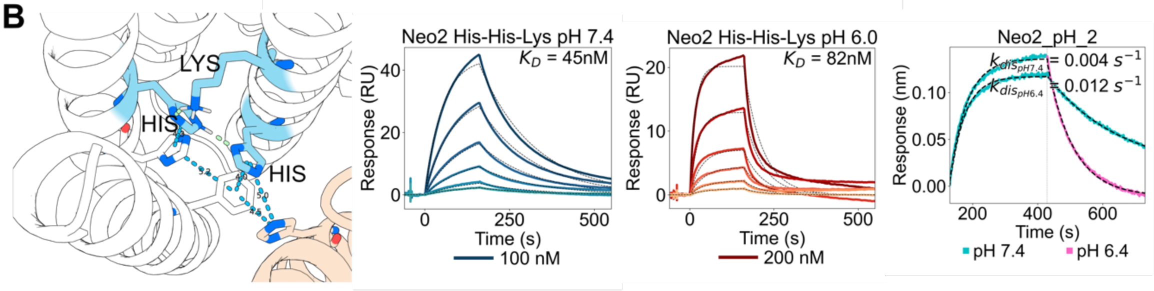


C

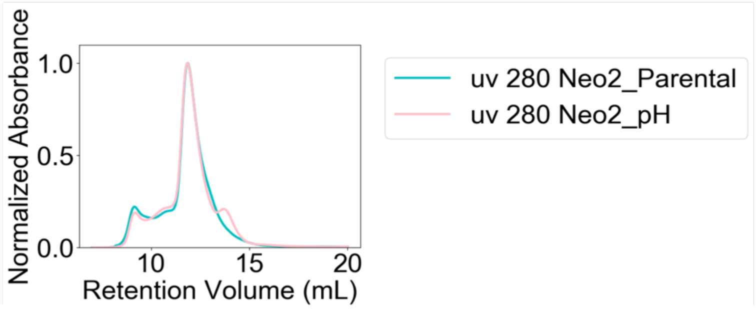


**Fig. S10. Characterizations of pH-sensitive Neo2 binders. A.** BLI curves of parental and pH-sensitive Neo2 binders. Association at pH 7.4 and dissociation at 6.4 showed faster dissociation at pH 6.4 for pH-sensitive binder. **B.** Design model and SPR curves of pH-sensitive Neo2 binder_2 with His-His-Lys network at pH 7.4 and 6.0. Association at pH 7.4 and dissociation at 6.4 showed faster dissociation at pH 6.4 for pH-sensitive binders. **C.** Size exclusion chromatography traces of pH-sensitive Neo2 binders.

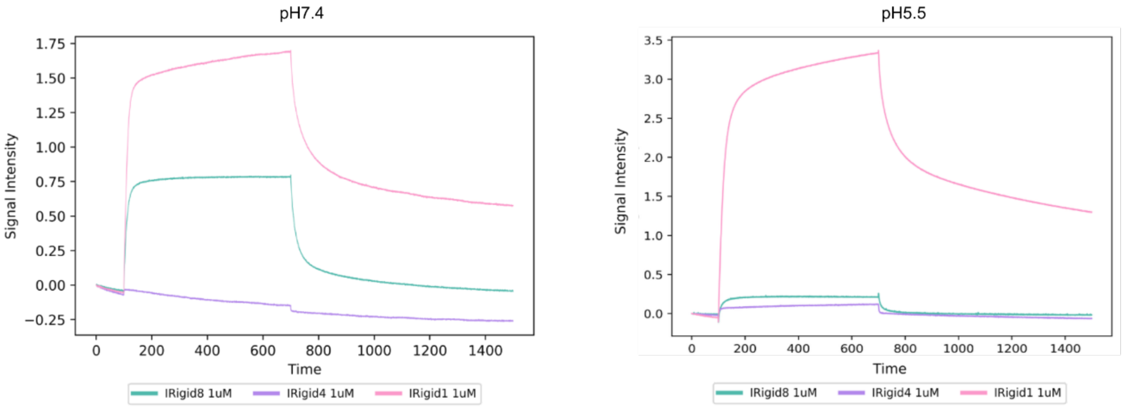


**Fig. S11**. **BLI binding curve of IGF2R Rigid (IRigid) binders at pH 7.4 and pH 5.5 to domain 11 of IGF2R.**

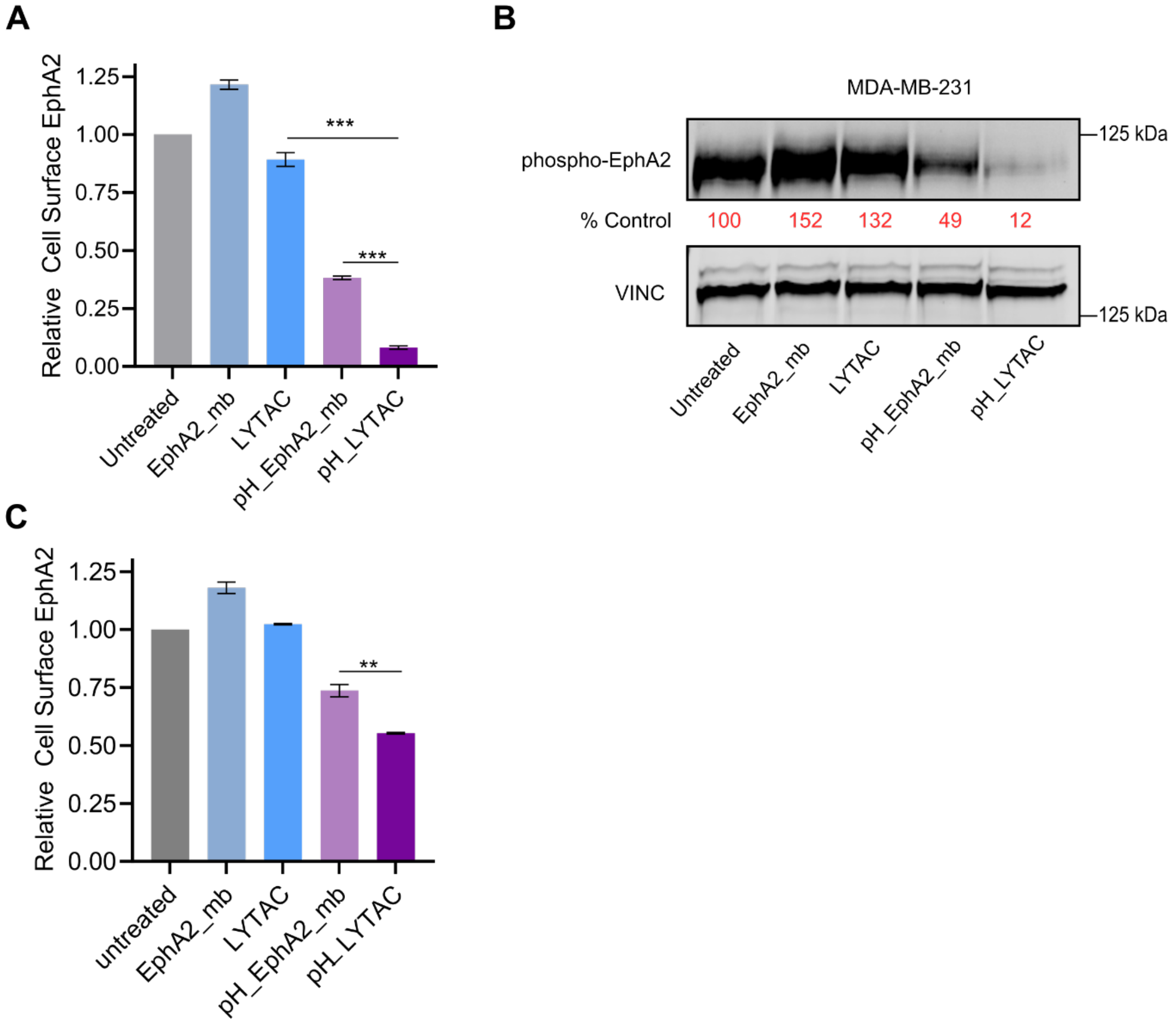


**Fig S12. Degradation of EphA2 in MDA-MB-231 cells. A.** Degradation of cell-surface EphA2 in MDA-MB-231 cells as determined by flow cytometry following 24h of 250 nM treatment. **B.** Degradation of phosphorylated EphA2 in MDA-MB-231 cells as determined by western blot following 24h of 250 nM treatment. **C.** Degradation of cell-surface EphA2 following a washout. MDA-MB-231 cells were treated with 250 nM of designed proteins for 24 h, then were washed out of extracellular treatment and incubated with fresh media for 6h.
